# Macrophage Polarizations in the Placenta and Lung are Associated with Bronchopulmonary Dysplasia

**DOI:** 10.1101/2024.01.26.577443

**Authors:** Karen K. Mestan, Abhineet Sharma, Sarah Lazar, Sonalisa Pandey, Mana M. Parast, Louise C. Laurent, Lawrence S. Prince, Debashis Sahoo

## Abstract

The intricate interplay between macrophage polarization and placenta vascular dysfunction has garnered increasing attention in the context of placental inflammatory diseases. This study delves into the complex relationship between macrophage polarization within the placenta and its potential impact on the development of vascular dysfunction and inflammatory conditions. The placenta, a crucial organ in fetal development, relies on a finely tuned balance of immune responses for proper functioning. Disruptions in this delicate equilibrium can lead to pathological conditions, including inflammatory diseases affecting the fetus and newborn infant.

We explored the interconnectedness between placental macrophage polarization and its relevance to lung macrophages, particularly in the context of early life lung development. Bronchopulmonary dysplasia (BPD), the most common chronic lung disease of prematurity, has been associated with abnormal immune responses, and understanding the role of macrophages in this context is pivotal. The investigation aims to shed light on how alterations in placental macrophage polarization may contribute to lung macrophage behavior and, consequently, influence the development of BPD. By unraveling the intricate mechanisms linking macrophage polarization, placental dysfunction and BPD, this research seeks to provide insights that could pave the way for targeted therapeutic interventions. The findings may offer novel perspectives on preventing and managing placental and lung-related pathologies, ultimately contributing to improved maternal and neonatal health outcomes.

## INTRODUCTION

The placenta plays a central role in supporting fetal development, orchestrating intricate processes crucial for a successful pregnancy. Macrophages, key components of the immune system, are integral players in maintaining the delicate balance required for proper placental function. Recent research has increasingly focused on understanding the nuanced phenomenon of macrophage polarization in a wide range of disease states, including pregnancy complications.^1-3^ However, little is known about macrophage polarization in the placenta and its impact on the developing fetus. Placental vascular dysfunction and inflammatory conditions pose significant threats to maternal and fetal health. Dysregulation of immune responses within the placenta can lead to adverse outcomes, including preterm birth and complications such as preeclampsia and intrauterine growth restriction.^4-6^ Unraveling the complexities of macrophage polarization in the context of placental pathology is essential for advancing our understanding of these conditions and exploring potential avenues for therapeutic intervention.

The connection between macrophage polarization and its influence on developing organ systems, particularly the lungs, is a subject of growing interest.^7-9^ However, there are no studies linking placental macrophage polarization and lung development. In the context of neonatal outcomes, bronchopulmonary dysplasia (BPD), a common developmental chronic lung disorder affecting premature infants, has been associated with aberrant immune responses, including lung macrophage activation and polarization. Investigating the crosstalk between macrophages in the placenta and the lungs holds promise for elucidating mechanisms underlying BPD development and progression.

Three recent studies on immune cell regulation in placental dysfunction and BPD provide rich and novel information on the transcriptomics of monocyte and macrophages in preterm birth. Sahoo, et al, described the gene expression profiles of lung macrophages in a cohort of preterm infants at risk for BPD.^10^ They measured changes in lung macrophage gene expression in premature infants at risk for BPD, and found higher inflammatory mediator expression with BPD, and with ex vivo response to LPS stimulation. In a subsequent report and different cohort, Sharma and colleagues reported that transcriptomic profiles in cord blood-derived fetal monocytes collected at birth reflect distinct placental dysfunction patterns from which the monocytes were drawn.^11^ Lastly, Windhorst, et al. conducted a similar study of cord blood monocyte subsets focusing on BPD outcomes.^12^ Collectively, the combined clinical, placental and immune cell data of these three pivotal studies can provide insight into the mechanisms by which placental function programs early lung development and later BPD outcomes.

Taken together, the 3 cohorts and supportive data surrounding them collectively suggest that cord blood monocytes are influenced by placental dysfunction, and that these altered monocyte progenitors—which are the precursors to early lung macrophages—are responsible for the placenta-lung crosstalk of BPD. An important gap remains whether the neonatal lung macrophages themselves correlate with prior placental dysfunction. Using a novel AI approach which can predict the function of tissue resident macrophages in any organ system, including the lung, we sought to understand how placental dysfunction drives changes in lung macrophages that resemble BPD. We hypothesized that the model would predict distinct patterns of placental-derived cord blood monocytes and infant lung macrophages that account for the BPD phenotype.

## STUDY DESIGN AND METHODS

### Description of Datasets

Three recently published datasets were included in the analysis. These 3 studies were conducted at 3 different sites. The published transcriptomic data on tracheal aspirate (GSE149490) and cord blood monocyte (GSE195727) samples were used to study the relationship between lung and placenta macrophages in SMaRT analysis as described below.

1. The cord blood monocyte dataset reported by Sharma, et al.^11^ This dataset consisted of bulk RNAseq data on monocyte subsets (classical, intermediate and non-classical) isolated from cord blood of preterm (N=59) and full-term (N=11) births at the time of delivery. The main objective was to compare monocyte subset gene expression according to preterm birth and placental inflammatory and vascular dysfunction (defined below).
2. The lung macrophage dataset reported by Sahoo, et al.^10^ This study included a total of 112 preterm infants born at <30 weeks gestation. All patients were intubated for mechanical ventilation due to respiratory distress syndrome. The initial sample for each patient was obtained within the first 24□h of life and subsequent samples were obtained weekly beginning on day 7 and continuing weekly if the patient remained intubated. Additional clinical data were linked to the datasets via medical chart reviews conducted by the research team at UCSD (SL and KM).
3. The cord blood monocyte dataset published by Windhorst, et al.^12^ This study enrolled 30 preterm infants born at <32 weeks gestational age with collection of cord blood for monocyte isolation into subsets (classical, intermediate, non-classical) as similarly isolated and sequenced by Sharma, et al.

### Determination of BPD

In all 3 studies, the NIH consensus definitions using endpoints of oxygen requirement and respiratory support at 36 weeks postmenstrual age was used to identify infants with BPD.^13^

### Categories of Placental Dysfunction

All patients in the monocyte study by Sharma, et al. had complete placental pathology data collected as part of the original study design. A comprehensive database of acute inflammatory, chronic inflammatory, maternal vascular and fetal vascular lesions was extracted from standardized pathology reports available by electronic medical records. The details of this approach have been previously published.^11, 14, 15^ There were no placental data collected in the original lung macrophage study by Sahoo, et al., however, the data were available in the electronic medical records for all infants born at our current institution (UCSD). Thus, we obtained all pathology reports of the infants born at UCSD under IRB-approved protocols and linked the placental data to the macrophage database. We applied the approach used in the cord blood monocyte study by Sharma, et al to assign primary placental domains for each of the 18 patients in the lung macrophage database (Figure 1C).

**Figure 1:**
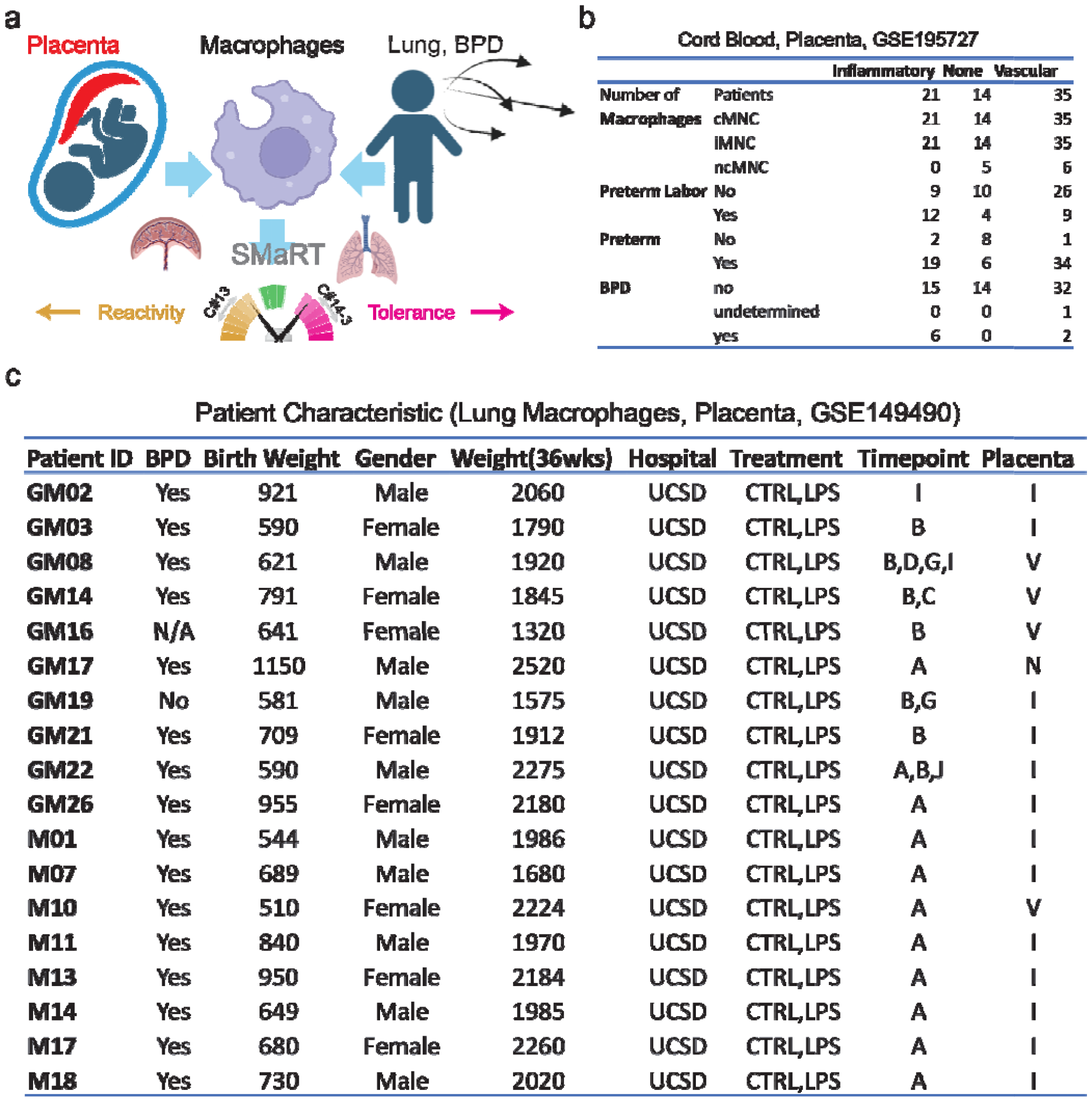
Study Design and Patient Characteristics. (a) Schematic experimental design that involve placenta phenotypes, cord blood monocytes, lung macrophages and BPD. (b) Patient characteristic of the cord blood study GSE195727. (c) Patient characteristic of the lung macrophage study GSE149490 with placenta phenotypes.

Each placenta was classified into 1 of 5 primary placental domains (acute inflammation, chronic inflammation, maternal vascular malperfusion, fetal vascular malperfusion, and none), based upon the presence and severity of gross and histologic lesions identified by the standard placental pathology exam and using the Amsterdam Workshop criteria.^16, 17^ Two broader categories of placental dysfunction were used in the predictive model: Placental inflammatory disease (PID) was defined as having either acute or chronic inflammation as the primary placental domain. Placental vascular disease (PVD) was defined as having either maternal or fetal vascular malperfusion as the primary placental domain.

#### SMaRT analysis

The Signatures of Macrophage Reactivity and Tolerance (SMaRT) offer a comprehensive quantitative and qualitative framework for evaluating macrophage polarization across various tissues and conditions.^18^ This study unveils a gene signature remarkably conserved across diverse tissues and conditions, comprising a set of 338 genes derived from a Boolean Implication Network model of macrophages.^18, 19^ This model effectively identifies macrophage polarization states at the single-cell level, encompassing a spectrum of physiological, tissue-specific, and disease contexts. Remarkably, this signature demonstrates robust associations with outcomes in several diseases, underscoring its potential as a valuable predictive tool. Boolean implication network has been used to identify universal biomarkers of macrophages earlier.^20^

The algorithm uses three clusters C#13, C#14, C#3 from the published macrophage network and uses composite scores of C#13, C#14-3, and C#13-14-3 to identify macrophage polarization states (See function getCls13, getCls14a3, getCls13a14a3, and orderData in github codebase BoNE/SMaRT/MacUtils.py and the outputs in BoNE/SMaRT/macrophage.ipynb). To compute the composite score, first the genes present in each cluster were normalized and averaged. Gene expression values were normalized according to a modified Z-score approach centered around StepMiner threshold (formula = (expr – SThr – 0.5)/3*stddev). Weighted linear combination of the averages from the clusters of a Boolean path was used to create a score for each sample. The weights along the path either monotonically increased or decreased to make the sample order consistent with the logical order based on Boolean Implication relationships. The samples were ordered based on the final weighted (−1 for C#13, 1 for C#14 and 2 for C#3) and linearly combined score. Performance is measured by computing ROC-AUC. Barplots show the ranking order of different sample types based on the composite scores of C#13, path C#14-3, or C#13-14-3. Violin plots shows the distribution of scores in different groups. P values are computed with Welch’s Two Sample t-test (unpaired, unequal variance (equal_var = False), and unequal sample size) parameters.

### Normalization of gene expression based on universal macrophage biomarker FCER1G

Since the number of macrophage varies between tissue samples, it is important to control for this variation during analysis of macrophage polarization. To start the normalization process, macrophage genes (such as FCER1G, etc.) is used to adjust the BoNE composite score. First, both the BoNE composite score and the macrophage gene expression are scaled for each sample type based on their dynamic range of expression values (min – max). For example, the dataset GSE220135, contains two sample types: noBPD, and BPD. Let’s take one sample from the BPD group (x, y) where x is the macrophage gene expression value and y is the original BoNE composite score. Bounding box for the BPD group demonstrates the range of values for both the BoNE composite score (S1) and the macrophage gene expression (S2). An average of BoNE composite scores and the macrophage gene expression is computed. The distance of (x, y) from the averages (S3, S4) is used to scale both values (x − S3*(S2 + 1)/(S1 + 1), y + S4*(S1 + 1)/(S2 + 1)). This process is repeated using noBPD samples using the averages. Linear regression is used to compute the trend between the transformed BoNE score and macrophage gene expression (y = mx + c). The trend is subtracted from the transformed BoNE score to compute the final normalized BoNE score (y = mx − c). Samples are now rank ordered based on the final normalized BoNE score to visualize the effect of normalization process.

### Statistical analyses

Gene signature is used to classify sample categories and the performance of the multi-class classification is measured by ROC-AUC (Receiver Operating Characteristics Area Under The Curve) values. A color-coded bar plot is combined with a density or violin + swarm plot to visualize the gene signature-based classification. All statistical tests were performed using R version 3.2.3 (2015-12-10). Standard t-tests were performed using python scipy.stats.ttest_ind package (version 0.19.0) with Welch’s Two Sample t-test (unpaired, unequal variance (equal_var = False), and unequal sample size) parameters. Multiple hypothesis corrections were performed by adjusting p values with statsmodels.stats.multitest.multipletests (fdr_bh: Benjamini/Hochberg principles). The results were independently validated with R statistical software (R version 3.6.1; 2019-07-05). Pathway analysis of gene lists were carried out via the Reactome database and algorithm.32 Reactome identifies signalling and metabolic molecules and organizes their relations into biological pathways and processes. Kaplan–Meier analysis is performed using lifelines python package version 0.14.6.

## RESULTS

### Study design to link placenta and lung macrophage polarization

We identified patients (**Fig 1a**) with placental vascular dysfunction and placenta inflammatory disease in previously published studies of cord blood monocytes (GSE195727) and lung macrophages (GSE149490).^10, 11^ Three different types of cord blood monocytes were profiled: classical (cMNC), intermediate (iMNC) and non-classical (ncMNC).^11^ To study macrophage polarization we employed the SMaRT (Signature of Macrophage Reactivity and Tolerance) model (**Fig 1a**). SMaRT model uses C#13 composite score to identify reactivity, C#14-3 to identify tolerance states, and C#13-14-3 to identify a simple overall summary for macrophage polarization.

In the cord blood monocyte study (GSE195727) we identified 21 patients with placental inflammatory disease (PID), 35 patients with placenta vascular disease (PVD) and 14 patients with none of these features (**Fig 1b**). Our hypothesis is that the monocytes from PID are more reactive compared to PVD. However, we anticipated that labor could be a confounding factor, based upon findings in the study by Sharma, et al. To study this further, we identified 25 patients with preterm labor: 12 patients have placental inflammatory disease, 9 patients have placenta vascular disease, and 4 patients with normal placenta.

To link placental dysfunction and lung macrophage polarization, we identified 13 patients with PID, 4 patients with PVD and 1 patient with none of these features (GSE149490, **Fig 1c**). Lung macrophage samples were taken from intubated patients within the first 24□h of life (A) and subsequent samples were obtained weekly beginning on day 7 (B) and continuing weekly (C-J) if the patient remained intubated.^10^ The lung macrophage samples were divided into control and LPS-treated conditions. 16 out of 18 patients had BPD.

### Preterm labor is associated with reactive macrophage polarization

Utilizing the SMaRT model based on C#13-14-3, cord blood monocytes underwent analysis to distinguish between reactive and tolerant states (**Fig 2a**). A cohort of preterm labor samples (n=54) exhibited a significant association with the reactive state (p = 0.000696). Notably, this correlation was evident across all monocyte populations, including classical, intermediate, and non-classical subsets. This observation aligns with the dynamic alterations in the maternal-fetal environment characterized by inflammation and tissue remodeling during preterm labor. In such an environment, monocytes are likely to undergo reprogramming, favoring the adoption of a reactive state.

**Figure 2:**
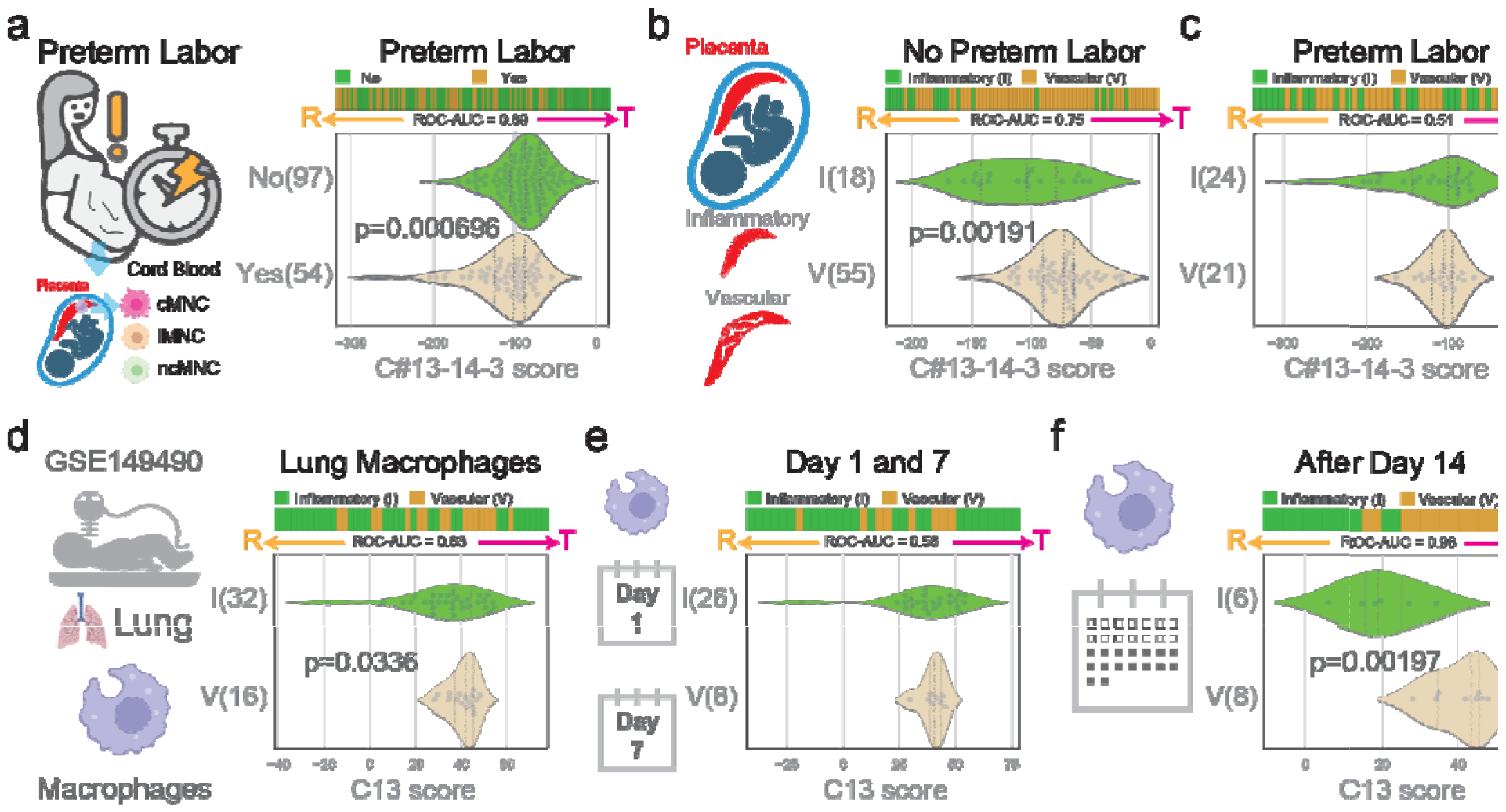
Vascular placenta is associated with tolerant macrophages in lung. (a) Preterm labor is associated with reactive cord blood monocytes. (b) Vascular placenta is associated with tolerant cord blood monocytes when there is no preterm labor. (c) No significant difference in cord blood monocytes polarization between Inflammatory and Vascular placenta when there is preterm labor. (d) Lung macrophage of babies born with vascular placenta retains tolerant phenotypes. (e) No significant difference between inflammatory and vascular placenta from lung macrophages in Day 1 and 7. (f) Lung macrophage after Day 14 of babies born with vascular placenta retains tolerant phenotypes.

### Placental vascular dysfunction is associated with tolerant macrophage polarization

Applying the SMaRT model based on C#13-14-3, we investigated the association between placental disease states and the polarization state of monocytes (**Fig 2b-c**). Notably, 18 samples from patients with PID displayed a significant reactivity score (low values) compared to the 55 samples from patients with PVD in the no preterm labor group (p = 0.00191). However, no significant difference was evident in the preterm labor group, possibly attributed to macrophage reprogramming during labor (**Fig 2a**). Anticipating a higher prevalence of reactive macrophages in patients with PID compared to those with PVD, particularly preceding preterm labor, underscores the dynamic nature of macrophage responses in these contexts.

### Macrophage polarization in lung is correlated with placenta

Examining lung macrophage samples through the C#13 SMART model revealed a reactive phenotype in patients with PID (**Fig 2d**, p = 0.0336), mirroring observations in cord blood from the placenta (**Fig 2b**). However, a detailed analysis indicated no significant difference in samples collected on the first day and day 7 (**Fig 2e**). Intriguingly, after day 14, samples from PID exhibited a significantly reactive state compared to Placenta Vascular Disease (PVD) (**Fig 2f**, p = 0.00197). Consistency in polarization phenotype across longitudinally collected samples every week suggests a post-birth reprogramming of macrophages, converging toward their associated placental phenotype after day 14. These findings highlight the dynamic nature of macrophage responses, emphasizing their potential adaptation and convergence in the postnatal period.

### BPD is associated with tolerant macrophage

The cord blood study GSE195727 exhibited a relatively small number of patients with BPD (Fig 3a). Analysis conducted using the C#13 SMaRT model indicates an association between BPD and tolerant states specifically within the intermediate monocyte subset (Fig 3b), contrasting with no such association observed in the classical monocyte subset (Fig 3c). This trend persisted across three additional publicly available datasets, including cord blood (GSE220135)^21^ and peripheral blood mononuclear cells (PBMC) (GSE125873, GSE108754),^22, 23^ consistently demonstrating that BPD is linked to tolerant states of macrophages (Fig 3d-f). These collective results underscore the robust and consistent association between BPD and the induction of tolerant macrophage states.

**Figure 3:**
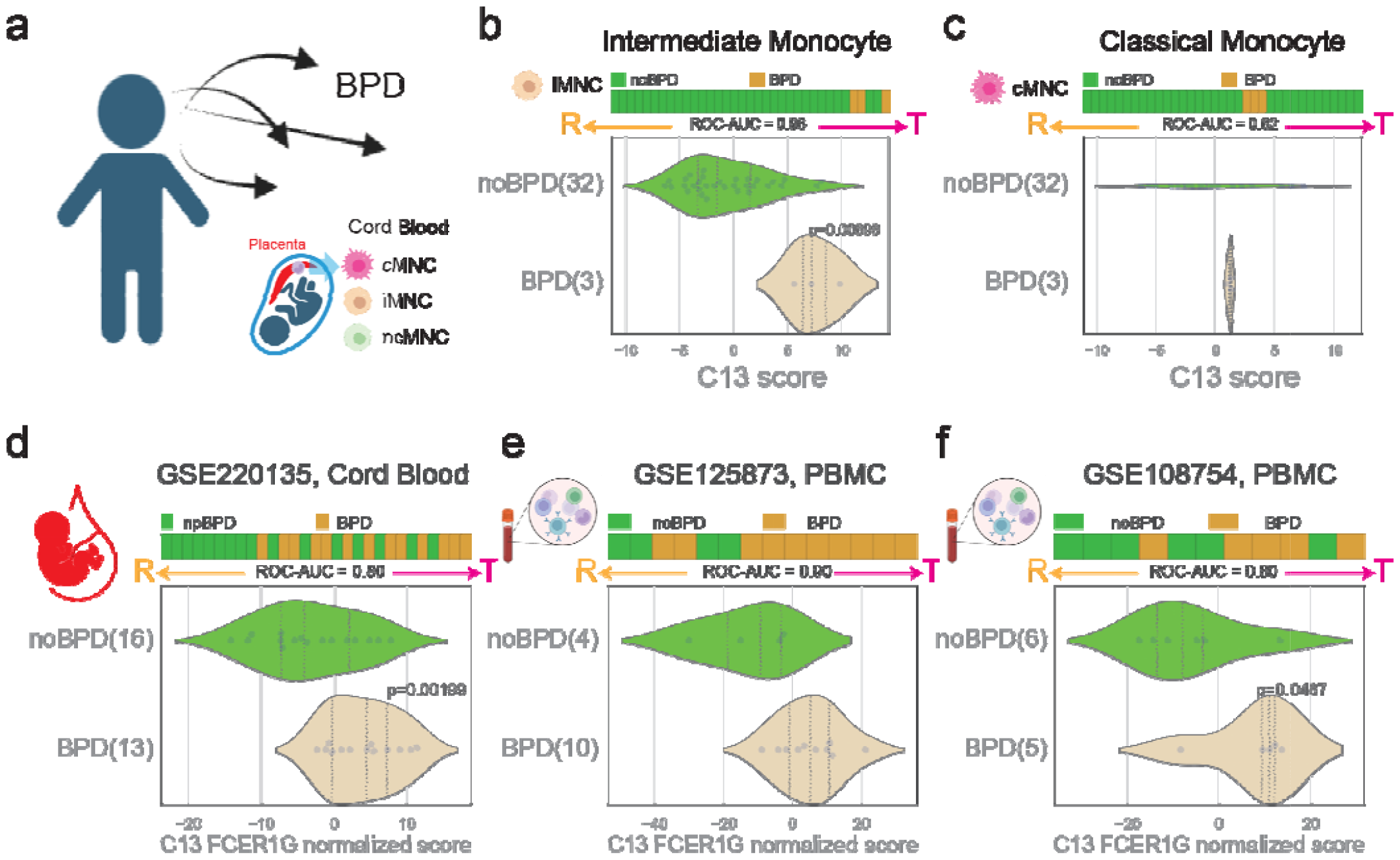
Tolerant monocyte/macrophages is associated with BPD. (a) Study design linking cord blood monocytes to BPD. (b) Tolerant polarization in intermediate cord blood monocyte is associated with BPD. (c) Tolerant polarization in classical cord blood monocyte is not significantly associated with BPD. (d) Tolerant monocyte polarization in cord blood (GSE220135) is significantly associated with BPD. (e) Tolerant monocyte polarization in PBMC (GSE125873) is significantly associated with BPD. (f) Tolerant monocyte polarization in PBMC (GSE108754) is significantly associated with BPD.

## DISCUSSION

In this study, we leveraged existing databases of clinical, placental and immune transcriptomic data coupled with novel analytic tools to evaluate macrophage polarization across placental and lung tissues with BPD outcomes. We found that preterm labor was associated with a reactive state of macrophages, and that this correlation is similar across monocyte subpopulations of classical, intermediate and non-classical subsets. We also found that, in the absence of preterm labor, immune cells from infants exposed to PID, characterized by histologic lesions of acute inflammation (chorioamnionitis, chorionic plate vasculitis, and/or funisitis), and chronic inflammation, was associated with a significant reactivity score (low values) compared to patients with PVD (maternal and/or fetal vascular malperfusion). Examining lung macrophage samples through the C#13 SMART model, we found a reactive phenotype in patients with PID, mirroring observations in cord blood from the placenta. While these associations were not found with lung macrophage samples collected in the first week of life, at later timepoints macrophages from patients exposed to PID exhibited a more prominent reactive state compared to PVD. Collectively, these findings suggest a dichotomy between inflammatory versus vascular-mediated processes in the developing lung that are regulated by immune cells programmed by distinct placental disease states.

Placental inflammatory lesions, such as chorioamnionitis (acute inflammatory lesions in the chorion and amnion), and funisitis (inflammatory infiltration in the umbilical cord and thus the fetus), are the most commonly associated placental findings associated with adverse neonatal outcomes.^24-27^ The association between placental inflammation and BPD has been variably and inconsistently reported, suggesting that the link between placental disease and fetal lung programming is more complex than previously understood.^28, 29^ More recent evidence has suggested that other pathophysiologic processes occurring *in utero* may lead to other forms of BPD mediated by mechanisms other than intrauterine inflammation.^30^ For example, placental vascular dysfunction, characterized by lesions of maternal and fetal vascular malperfusion in which abnormal trophoblast implantation, failed spiral artery remodeling, decidual and/or fetal vasculopathies and other events lead to chronic fetal hypoxia. The importance of placental vascular findings on neonatal outcome has recently been highlighted by robust associations between placental vascular disease and BPD with pulmonary hypertension—a severe form of BPD associated with higher morbidity and mortality.^14, 31^ The findings of this study provide new mechanistic insights into how distinct forms of placental dysfunction lead to distinct endo-phenotypes of BPD.

Placental health is intricately linked to the balance of macrophage populations, with PID often characterized by the presence of reactive macrophages. In contrast, PVD exhibits a different macrophage profile. These distinct macrophage populations within the placenta have the potential to exert a lasting impact on the polarization of macrophages in other organs, particularly the lungs.^32-34^ The process of labor and birth introduces a significant shift in the immune milieu, reprogramming macrophages to accommodate the dynamic changes associated with parturition. This reprogramming, however, tends to blur the differences between macrophage states associated with various disease conditions. Despite this, the characterization of placental macrophages before the onset of labor becomes a crucial snapshot in understanding the potential long-term programming of macrophages.

The intricate relationship between labor and monocyte/macrophage polarization is a fascinating yet understudied area of study within reproductive immunology.^35-37^ Monocytes, as key components of the innate immune system, play a crucial role in the dynamic processes associated with labor, contributing to both the initiation and resolution of this complex physiological event. During labor, the uterine environment undergoes significant changes, marked by inflammation and tissue remodeling. Macrophages within the uterine tissues exhibit a shift in their polarization states, transitioning between reactive/pro-inflammatory (M1) and tolerant/anti-inflammatory (M2) phenotypes. This dynamic polarization is orchestrated to facilitate the various stages of labor, including cervical ripening, uterine contractions, and postpartum tissue repair.

By examining the macrophage landscape in the placenta before labor, we may gain insights into the initial conditions that influence the trajectory of macrophage polarization in distant organs, such as the lungs. This long-term influence on lung macrophages is particularly relevant in the context of BPD. Understanding how cord blood-derived placental-fetal monocytes, specifically those in an inflammatory state, contribute to the programming of lung macrophages provides a unique perspective on the etiology of BPD. The consistent reflection of pre-labor placental monocytes in the programming of lung macrophages suggests a potential link between placental health and the development of respiratory conditions in neonates. Further research in this direction may uncover novel therapeutic strategies aimed at modulating macrophage behavior to mitigate the risk of BPD and improve neonatal outcomes.

Important limitations of this analysis include the differences in cohort size and differences in the primary focus of each study that required retrospective collection of placental data in the lung macrophage study. While placental pathology is infrequently the focus in longitudinal cohort studies involving the developing neonate, our findings confirm that the placenta should play a pivotal role in mechanistic studies linking early biomarkers and outcomes. New AI technologies are emerging that can link vast amounts of transcriptomic and other data across different cohorts and over time. These approaches provide exciting new possibilities to elucidate mechanisms of complex diseases such as BPD that require longitudinal study of immune and vascular processes across the lifespan.

In conclusion, the completed analysis serves as a novel model for studying BPD and its endotypes. More comprehensive investigation of the intricate relationships between macrophage polarization and BPD mediated by placental dysfunction are needed. By shedding light on these interconnected processes, these discoveries will contribute valuable insights that may inform novel therapeutic strategies for mitigating the impact of placental and lung-related pathologies on maternal and neonatal health.

## Contributors

Conceptualization: DS, KKM

Methodology: DS, KKM, AS, SL, LSP, MMP

Investigation: DS, KKM, AS, SL, SP, LSP, LCL, MMP

Visualization: DS, KKM, SP

Funding acquisition: DS, KKM, LCL, LSP, MMP

Project administration: DS, KKM

Supervision: DS, KKM

Writing – original draft: DS, KKM

Writing – review & editing: DS, KKM, AS, SL, SP, LSP, LCL, MMP

DS, and KKM have accessed and verified the underlying data. All authors read and approved the final version of the manuscript.

## Data Sharing

All data are available in the main text or the supplementary materials. The codes are available in https://github.com/sahoo00/BoNE.

## Acknowledgements

This work was supported by the National Institutes for Health (NIH) grant R01-AI155696 (to DS), R01HL139798 (to KKM), U01HL126494 (to LCL), UG3CA241687 (to LCL), and R01HD089537 (to MMP). Other sources of support include: R01-GM138385 (to DS), and UG3TR003355 (to DS). Images in the figures were created with BioRender.com.

